# Coordinated regulation of WNT/β-catenin, c-Met, and Integrin signalling pathways by miR-193b controls triple negative breast cancer metastatic traits

**DOI:** 10.1101/2021.05.10.443372

**Authors:** Chiara Giacomelli, Janine Jung, Astrid Wachter, Susanne Ibing, Rainer Will, Stefan Uhlmann, Heiko Mannsperger, Özgür Sahin, Yosef Yarden, Tim Beißbarth, Ulrike Korf, Cindy Körner, Stefan Wiemann

**Affiliations:** Division of Molecular Genome Analysis, German Cancer Research Center (DKFZ), Heidelberg, Germany; Medical Bioinformatics, University Medical Center Göttingen, Göttingen, Germany; Division of Applied Bioinformatics, German Cancer Research Center (DKFZ), Heidelberg, Germany; Genomics and Proteomics Core Facilities, German Cancer Research Center (DKFZ), Heidelberg, Germany; Biological Regulation, Weizmann Institute of Science, Rehovot, Israel

## Abstract

**Background:** Triple Negative Breast Cancer (TNBC) is the most aggressive subtype of Breast Cancer (BC). Treatment options for TNBC patients are limited and further insights into disease aetiology are needed to develop better therapeutic approaches. microRNAs’ ability to regulate multiple targets could hold a promising discovery approach to pathways relevant for TNBC aggressiveness. Thus, we address the role of miRNAs in controlling signalling pathways and phenotypes relevant to the biology of TNBC.

**Methods:** To identify miRNAs regulating WNT/β-catenin, c-Met, and integrin signalling pathways, we performed a high-throughput targeted proteomic approach, investigating the effect of 800 miRNAs on the expression of 62 proteins in the MDA-MB-231 TNBC cell line. We then developed a novel network analysis, Pathway Coregulatory (PC) score, to detect miRNAs regulating the three pathways. Using *in vitro* assays for cell growth, migration, apoptosis, and stem-cell content, we validated the function of candidate miRNAs. Bioinformatic analyses using BC patients’ datasets were employed to assess expression of miRNAs as well as their pathological relevance in TNBC patients.

**Results:** We identified six candidate miRNAs coordinately regulating the three signalling pathways. Quantifying cell growth of three TNBC cell lines upon miRNA gain-of-function experiments, we characterised miR-193b as a strong and consistent repressor of this phenotype. Importantly, the effects of miR-193b were stronger than chemical inhibition of the individual pathways. We further demonstrated that miR-193b induced apoptosis, repressed migration, and regulated stem-cell markers in MDA-MB-231 cells. Furthermore, miR-193b expression was the lowest in patients classified as TNBC or Basal compared to other subtypes when classified by PAM50 signatures. Gene Set Enrichment Analysis showed that miR-193b expression was significantly associated with reduced activity of of WNT/β-catenin and c-Met signalling pathways in TNBC patients.

**Conclusions:** Integrating miRNA-mediated effects and protein functions on networks, we show that miRNAs predominantly act in a coordinated fashion to activate or repress signalling pathways responsible for metastatic traits in TNBC. We further demonstrate that our top candidate, miR-193b, regulates these phenotypes to an extent stronger than individual pathway inhibition, thus proving that its effect on TNBC aggressiveness is mediated by repressing multiple interconnected pathways.

## BACKGROUND

Triple negative breast cancer (TNBC) is a heterogeneous subtype of breast cancer, histologically characterised by the absence of expression of oestrogen- (ER), progesterone- (PR), or HER2 receptor expression. Compared to other breast cancer subtypes, TNBC displays the lowest 5-year survival rates, regardless of the stage at diagnosis (Surveillance, Epidemiology, and End Results - SEER 2019). Additionally, TNBC patients’ 5-year survival dramatically decreases to 65% and 12.2% if the disease had already spread to regional lymph nodes or at distal sites at the time of diagnosis, respectively (SEER 2019). Metastatic recurrence has remained the main cause of cancer-related deaths for all breast cancer patients (1) and thus represents a major challenge for TNBC patients. Indeed, they present the highest percentages in both local and distant recurrences, with metastases more common in brain and lungs (2). As well, median duration of survival with distant metastasis is the lowest for TNBC (0.5 years) compared to other subtypes (2.2 for LumA, 1.6 LumB, 0.7 Her2+) (3).

The intrinsic heterogeneity of TNBC tumours is a double-edged sword, concomitantly underlying unpredictable differences in response to chemotherapeutic treatments while also presenting itself as potential source of therapeutic vulnerabilities to explore. For the majority of TNBC patients the only viable treatment option is chemotherapy, with responses ranging from pathological complete response (pCR) associated with high rates of survival (30 to 40% of patients), to residual disease after neoadjuvant treatment, a prognostic factor of extremely poor survival (4). More recently, four subtypes of TNBC were identified, Basal-Like1 and 2 (BL1 and BL2), mesenchymal (M), and Luminal Androgen Receptor (LAR). BL2 patients have the lowest probability of reaching a pCR among all TNBC subtypes, and the lowest distant relapse free survival (2). Thus, TNBC as a heterogeneous disease and the BL2 subtype specifically require deeper biological investigations to fully understand the pathological mechanisms, which underlie its clinical aggressiveness, as well as to identify viable novel therapeutic avenues.

The prime function of microRNAs (miRNAs) is to negatively regulate the expression of genes at their post-transcriptional level by interacting with the 3’ untranslated region (UTR) of respective targets. The extent of this regulation has been characterized both at the transcriptomic and proteomic levels, indicating that while regulating a multitude of targets, this happens in a mild fashion. Indeed, various studies have used gain-of-function or loss-of-function miRNA experiments that showed an effect ranging between -0.3 log2FC (5) and +0.15 log2FC (6), respectively. In more recent years, the scientific community came to appreciate that miRNAs functionally relevant for specific phenotypes regulate multiple targets within the same signalling cascade (7). Indeed, a high throughput screening (HTS) at the proteomic level identified miR-193a, miR-124, and miR-147 as regulators of proliferation dependent on their function on the EGFR signalling pathway (8). Additionally, members of the miR-200 family were characterized to cumulatively affect proteins involved in actin cytoskeleton remodelling, regulating invasion and invadopodia formation (9). In vivo, the combinatorial role of miRNAs belonging to the miR-17-92 cluster has been dissected, identifying how each of them contributes to specific phenotypes identified by the depletion of the cluster as a whole (10). As well, in adult mouse neurons, miR-128 was identified as a decisive regulator of neuronal excitability, due to its ability to control the expression of various ion channels and ERK2 signalling (11). A recent review has revisited in depth all these phenotypes and network functions of miRNAs showing how they might be additionally integrated in feed-forward and feedback networks, providing insights into the effects that miRNAs have in the context of cancer (7).

Due to their dose-sensitivity, biological pathways require fine-tuned control of the signalling cascade. In these contexts, miRNAs may become pivotal regulators, thanks to their ability to direct the expression of multiple targets (12). Thus, in this study we aimed to identify miRNAs with a functional relevance in TNBC, mediating a coordinated regulation of signalling pathways. We focused on the WNT/β-catenin, c-Met, and Integrin signalling pathways due to their enrichment in the BL2 subtype, which is characterised by worse clinical features (13). To address the global effects of miRNAs on these pathways, we performed a targeted quantification of proteins upon miRNA gain-of-function in MDA-MB-231 cells, a model of BL2 TNBC. Subsequently, we developed a novel network analysis integrating the effect of miRNAs on proteins’ expression with the function of the same proteins on the pathways of interest, an essential information frequently overlooked in network analysis approaches. We further characterised miR-193b as a new strong repressor of all three pathways in TNBC, validating its regulatory effects in gene expression data derived from patients’ datasets.

## MATERIALS AND METHODS

### Cell culture

The human triple negative breast cancer cell lines MDA-MB-231 (Cellosaurus:CVCL_0062) and HCC-1806 (CVCL_1258) were obtained from ATCC (Manassas, VA, USA). SUM-159 (CVCL_5423) cells were a kind gift from Andreas Trumpp (DKFZ, Heidelberg, Germany). All cell lines were authenticated using Multiplex Cell Authentication by Multiplexion (Heidelberg, Germany) as previously described (14). The SNP profiles matched the expected ones. All cell lines were routinely tested for potential contamination with mycoplasma. MDA-MB-231 cells were cultured in Leibovitz-L15 medium (Gibco, Thermo Fisher Scientific) supplemented with 10% foetal calf serum (Gibco) and 3 g/lt of sodium bicarbonate. HCC-1806 cells were cultured in RPMI-1640 (Gibco), supplemented with 10% foetal calf serum (Gibco). SUM-159 cells were cultured in Ham’s F12 (Gibco), supplemented with 5% foetal calf serum (Gibco), 10mM Hepes (Gibco), 10 ug/ml Hydrocortysone (BRAND), and 5 ug/ml Insulin (BRAND). All cell lines were cultured in incubators maintained at 37C and 5% CO_2_.

### microRNA gain-of-function experiments

The mimic overexpression screening and cell pellet retrieval were performed as previously described (8). All additional transient transfections were performed with Lipofectamine2000 (Invitrogen, CA, USA) according to manufacturer’s instruction. miRNA mimics and respective negative controls, siRNA and respective siRNA control were purchased from Dharmacon (GE Healthcare) and used at a final concentration of 25nM. Unless otherwise stated, the miRNA negative controls used were miRIDIAN microRNA Mimic Negative Control #1 and #2.

### Ectopic activation and inhibition of signalling pathways

The WNT/β-catenin pathway was stimulated with mouse recombinant WNT3a (Peprotech, NJ, USA) at a final concentration of 100 ng/ml. β-catenin transcriptional activity was inhibited by treating cells with iCRT14 (Santa Cruz, CA, USA) at a final concentration of 10 µM. c-Met signalling was stimulated with recombinant human HGF (R&D Systems) at a final concentration of 75 nM, whilst it was inhibited with Capmatinib (Biozol Diagnostica, Eching, Germany) at a final concentration of 2 nM. c-Met and EGFR signalling were concomitantly stimulated with recombinant human EGF (Corning, NY, USA) at a final concentration of 20 nM. Downstream signalling was inhibited with Erlotinib at a final concentration of 5 µM. Recombinant WNT3a, HGF, and EGF were diluted in 0.1% BSA in PBS, which was therefore used as a control in all experiments and is indicated with the “unstimulated” label in respective figures. iCRT-14 and Capmatinib were diluted in DMSO, whilst Erlotinib was diluted in PBS. Thus, the respective vehicle controls (veh. ctrl) were used in the experiments.

### RPPA

RPPA was performed as previously described (15). Briefly, protein lysates harvested from miRNA-overexpressing MDA-MB-231 cell-pellets were thawed and printed in technical triplicates on nitrocellulose coated glass slides (Oncyte Avid, Grace-Biolabs) using a contact spotter (Aushon BioSystems). Lysates were separated into four groups for spotting. Each of them included appropriate dilution controls for downstream analysis as well as samples transfected with miRNA mimic controls 1 and 2, employed for differential expression analysis (see RPPA HTS data analysis paragraph). Antibody validation for the RPPA screening was performed as previously described (16). Supplementary table 1 lists all antibodies used in this study. Unless otherwise noted in Supplementary table 1, the antibodies were incubated at 1:300 dilution in Blocking buffer. After four washes in Washing buffer, primary antibodies were detected using Alexa Fluor 680 F(ab’)2 fragments of goat anti-mouse immunoglobulin G (IgG) or anti-rabbit IgG (Life Technologies) diluted at 1:8,000 in Blocking buffer. Images were acquired at 700 nm wavelength and with 21 µm resolution using an Odyssey scanner (LI-COR, NE, USA). Every nine slides, one was reserved for total protein content analysis by staining with the Fast Green FCF method (17). Signal intensities were quantified using the GenePix Pro software v.7 (Molecular Devices, CA, USA).

### RPPA HTS data analysis

Signal intensities were processed using the R package RPPanalyzer (v. 1.4.3) (18) for quality control and total protein content normalization. Data quality was assessed by i) checking target specific signals in comparison to their corresponding blank values of the serially diluted control samples and by ii) comparing target measurement signals against blank signals. Spot-wise normalization to the total protein concentration was performed based on the Fast Green FCF method (17). Potential block effects were removed by shifting the median value of each block to the overall median. The R package ‘limma’ (version 3.26.9) (19) was used to identify miRNAs causing a differential expression of proteins. Within each transfection round, the signal intensities of the miRNA overexpression samples were tested against the two mimic negative control values. Specifically, the comparison was performed between miRNA-transfected samples in two biological replicates against mimic control-transfected samples in two biological replicates of two distinct negative controls. Multiple testing correction was performed with the Benjamini-Hochberg method (20). For downstream analyses and data plotting, only miRNAs causing at least one statistically significant alteration across the dataset were considered, leading to a final data table and corresponding heatmap of 722 miRNAs by 62 proteins. Statistical significance threshold: q-value ≤ 0.001. Data analyses were performed in R version 3.2.2.

### Enrichment analyses

Two databases were assessed to retrieve miRNA-target predicted relationships: TargetScanHuman (v.7) (21) and MicroCosm Targets (previously known as miRBase::Targets) (22). TargetScanHuman database information for conserved and non-conserved targets was individually analysed. Fisher’s exact test was used for enrichment testing, individually addressing downregulated miRNA-target pairs and upregulated miRNA-target pairs. Differential protein expression was considered significant below a threshold of q-value ≤ 0.001.

### Pathway Coregulatory score analysis

For pathway analysis, protein effects caused by miRNAs (q-value ≤ 0.001) were combined with the respective regulatory protein function in the pathway (either activator or repressor) following the rules illustrated in Fig. 1C. Briefly, downregulation of an activator protein in the pathway resulted in a negative pathway effect, and downregulation of a repressor protein in the pathway resulted in a positive pathway effect. The opposite was considered if the miRNA was causing the upregulation of protein expression, Supplementary table 4 describes the list of targets and the respective biological effect on associated pathway(s). A pathway coregulation (PC) score was defined for each miRNA as the sum of all measured miRNA-mediated effects on the pathway, weighted by the number of measured proteins in the pathway. Permutation testing was performed to assess the miRNA-wise probability distribution of PC scores by 10,000x resampling the significant miRNA-protein interactions for each protein. PC scores were considered significant based on a 5% FDR. Supplementary file 1 contains the R code in html format employed for the bootstrap analysis.

**Figure 1.**
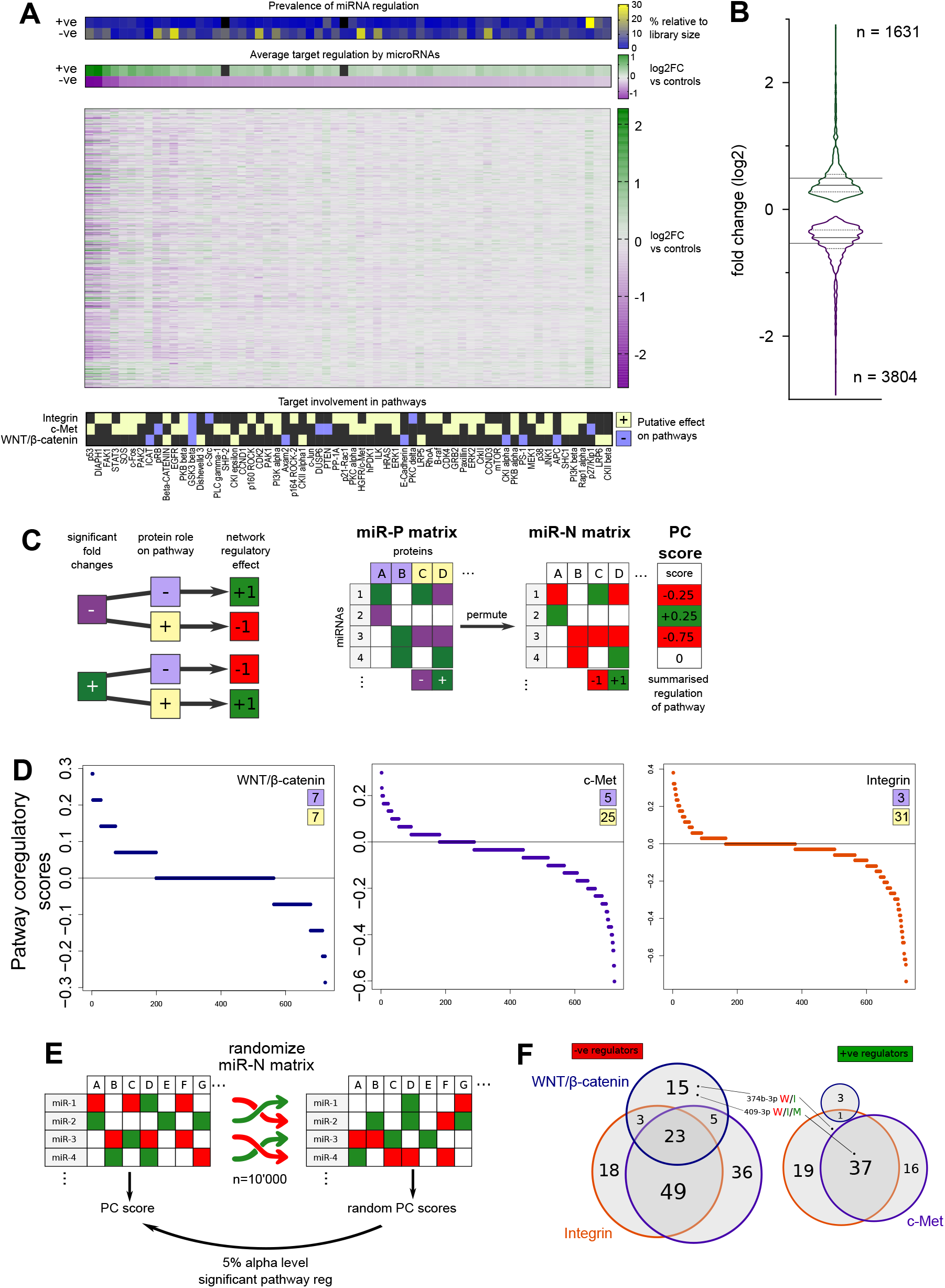
miRNAs coordinately regulates signalling pathways despite mildly regulating individual targets. **A** and **B** – MDA-MB-231 cells were transfected with individual miRNAs from a library of 800 representing the global miRNome. 48hrs post-transfection, total protein lysates were harvested and the expression of 62 target proteins was assessed by means of Reverse Phase Protein Assay (RPPA). After normalization for total protein content with FCF, the effect of miRNAs on target proteins was quantified by limma test. P-values were corrected for multiple testing with Benjamini-Hochberg method. Tabular results are available in Supplementary table 2. A. Effects of miRNAs on the 62 probed targets are represented with a heatmap of all fold changes compared to negative controls in log2 scale (log2FC). Each column is a target, each row a microRNA. Only miRNAs which caused at least one statistically significant interaction across the entire dataset were plotted, leading to 722 rows. The upper rug represents the prevalence of miRNA regulation for each target, weighted for the library size, separating the number of miRNAs significantly positively (+ve) or negatively (-ne) regulate each target. The second rug represents the average of statistically significant (q-value ≤ 0.001) regulation of each target, separating positive and negative regulations. The lower rugs represent from which pathway(s) of origin were the targets derived from as well as the putative effect of the target on the pathway. B. The regulatory activity of miRNAs is summarised in violin plot of all statistically significant (q ≤0.001) log2FC in protein expression, maintaining separated positive and negative regulations. Full and dotted lines in the violins respectively represent the medians and the quartiles of the distributions. The horizontal lines in the plot represent the averages. Number of statistically significant upregulations and downregulations are written in the respective area of the plot. C. Principles of the computation of PC scores. A set of rules is exploited to mathematically transform the effect of a miRNA on a single target protein into a Pathway Coregulatory (PC) effect. The score is designed to integrate the function of the assayed protein on the signalling pathway: a miRNA might negatively or positively regulate (dark purple and green, respectively) the expression of a target with repressive or activating function (lilac and yellow, respectively). Depending on the combination of these two factors, the effect on the pathway is positive or negative (bright green or red, respectively). The cumulative effect of a miRNA on the pathway is then summarized in a PC score classifying each miRNA as activator or repressor of a pathway. D. The distribution of PC scores shows the global effects of miRNAs on the pathways The three individual pathways probed are displayed: WNT-beta-catenin (left), c-Met (middle), and Integrin signalling (right). In the upper corner of each graph numbers indicate the number of putative repressing or activating proteins probed associated to the pathway. miRNAs are ranked within each pathway by highest to lowest computed PC score. E. Principles of statistical testing of the computed PC scores. The miR-N matrix used to calculate the PC scores is randomized 10’000, for each a miRNA-specific random PC score is computed. Then, the experimental PC score is tested against the randomly generated ones. An experimental PC score is considered significant with a 5% alpha level. F. Venn diagrams display the number of miRNAs with a statistically significant effect on pathway regulation after randomization test. The left group indicates the number of miRNAs repressing the pathways, while on the right the number of miRNAs activating the pathways.

### Generation of isogenic recombinant cell lines for WNT pathway reporter assay

MDA-MB-231 were generated as described (23). Briefly, a mammalian expression vector (pPAR3) containing a Flp recombinase target site N-terminally fused to EGFP under control of an elongation factor 1-alpha (*eEF1a1*) promoter and a neomycin selection marker, was stably integrated in the genome of MDA-MB-231 cells. Neomycin resistant and EGFP positive clones were isolated and validated for single-copy integration of the FRT site by Southern blotting. Functionality of the MDA-MB-231-pPAR3 acceptor cell line was verified using Flp-mediated recombination with either a Doxcylin inducible hcRED expression vector for visual testing or a red firefly expression vector for quantitative expression analysis. The validated MDA-MB-231-pPAR3 acceptor cell line served a platform for the generation of isogenic variants.

For generation of MDA-MB-231-pPAR3 WNT/β-catenin-Pathway reporter cell lines, a dual reporter vector containing a promoter-less FRT reporter cassette with TCF/LEF responsive elements followed by a cassette for normalization was flipped into the MDA-MB-231-pPAR3 acceptor cell line by co-transfection with a Flp recombinase expression vector (pOG44 / Invitrogen). The WNT reporter cassette consists of a RNA polymerase II transcriptional pause signal from the human hemoglobin subunit alpha 2 gene (*HBA2*) followed by 6 repeats of the TCF/LEF transcriptional response element (AGATCAAAGGGGGTA) joined to a minimal TATA-box promoter and destabilized firefly luciferase reporter (Qiagen, CCS-018L). The cassette for normalization contains a SV40 promoter driving the renilla luciferase open reading frame, which allows dual measurement of both luciferases. After selection for hygromycin resistance (expression vector) and loss of EGFP expression (positive integration), single colonies were picked and analysed in in the presence of recombinant WNT with dual luciferase assays for WNT / luciferase responsiveness and renilla luciferase expression.

### WNT pathway reporter assay

Three clones of isogenic MDA-MB-231-pPAR3 WNT/β-catenin reporter cell lines were plated at the density of 10’000 cells/well in white flat bottom 96-well plates. The subsequent day, cells were transfected with indicated miRNA mimics or controls. Alternatively, cells were treated with iCRT14. The next day, cells were stimulated with recombinant WNT3a and 18hrs later the dual luciferase activities were assayed using a luminometer (Tecan, Männedorf, Switzerland). The median of six technical replicates was used to calculate the ratio over the control. Ratios for three independent clones were averaged and used for statistical testing (two-tailed, unpaired t-test). Statistical testing was performed using GraphPad Prism v9.

### Proliferation and apoptosis assays

Cell lines were plated in black 96-well plates with clear bottom at the indicated densities, based on their respective growth rates: MDA-MB-231 cells at 5’000 cells/well, SUM-159 at 1’700 cells/well, and HCC-1806 at 2’000 cells/well. The next day, cells were transfected with indicated miRNA mimics or negative controls using Lipofectamine 2000. Alternatively, cells were treated with iCRT14, Capmatinib, or Erlotinib. Pathway stimulation was performed concomitantly with chemical inhibition or, for miRNA transfection, 5 hrs post-transfection upon medium change.

72 hrs post-treatment, nuclei of cells were stained with Hoechst 33342 (Life Technologies) at a final concentration of 20 µM for 30 minutes at 37C. Subsequently, they were imaged using a Molecular Devices Microscope IXM XLS with 4x S Fluor objective. To assay for apoptotic cells, cells were additionally stained with Propidium Iodide (PI, Life Technologies) at 0.2 ng/mL in addition to Hoechst staining. PI was added 5 minutes prior to imaging using a Molecular Devices Microscope IXM XLS with 4x S Fluor objective. The percentage of apoptotic cells was counted by normalizing the PI-positive nuclei to the number of total nuclei (stained with Hoechst).

Image analysis was performed with built-in software. Six technical replicates were performed for each experiment, the mean of the technical replicates was used to calculate the ratio of treatment over control. Ratios of three independent biological replicates were used for statistics (two-tailed, unpaired t-test). Statistical testing was performed using GraphPad Prism v9.

### Migration assay

MDA-MB-231 cells were plated into clear 6-well plates (Greiner Bio-One) at 400’000 cells/well. The next day, they were either transfected with miR-193b or negative control #2. Alternatively, they were treated with iCRT14 or DMSO control. Two days after transfection, the cells were starved for 18h with serum-free medium. Subsequently, 200’000 cells were reseeded in serum-free conditions into the upper compartment of 6.5 mm transwell inserts with 5.0 µm pores (Corning), while medium with 10% FCS in the lower compartment was used as chemoattractant. To mimic the inhibitory effect caused by miR-193b overexpression in cells, iCRT14 or DMSO was added both during starvation and upper chamber reseeding, but not in the lower chamber. Overall, reseeding was performed 72 hrs post-transfection or treatment and migration readout was performed 20h after reseeding. Inserts were briefly washed with PBS, then cells that had remained within the upper chamber were removed with a cotton swab. Cells that had migrated through the pores were fixed with 4% PFA (prepared from 16% Formaldehyde (w/v), Thermo Fisher Scientific) for 15 minutes at the lower side of the insert membrane and stained with 20 µM Hoechst 33342 (Life Technologies) for 30 minutes. Cells were imaged using a Molecular Devices Microscope IXM XLS with 4x S Fluor objective and quantified with built-in software. The mean of three technical replicates was calculated and normalized to a seeding control to account for differences in reseeded cell numbers. Subsequently, a ratio to control condition was calculated and the average for three biological replicates was used for statistics (two-tailed, unpaired t-test). Statistical testing was performed using GraphPad Prism v9.

### FACS analysis of CD24 and CD44 surface expression markers

MDA-MB-231 cells were plated in 6-well plates at a density of 250’000 cells/well. The next day cells were transfected with miR-193b or mimic negative control #2. Alternatively, cells were treated with iCRT14 or DMSO as control. Four days after transfection, cells were detached with Cell Dissociation Buffer, enzyme-free (GIBCO) and stained for CD-44 and CD-24 surface markers with PE- and APC-conjugated antibodies, respectively (Biolegend). Unstained and isotype controls for APC and PE (Biolegend) were used as controls to gate positive cells. Stained samples were immediately analysed on a FACSCanto II (BD Biosciences). The results were analysed using FACSDiva software (v8, BD Biosciences). Cell percentages from six independent biological replicates with two technical replicates each were used for statistical testing (two-tailed, paired t-test). Statistical testing was performed using GraphPad Prism v9.

### Patient data analysis from TCGA and Metabric BRCA datasets

miRNA isoform expression quantification data of patients from the TCGA cohort was generated by the TCGA Research Network (https://www.cancer.gov/tcga). The workflow to process the data was based on the British Columbia Genome Sciences Centre miRNA Profiling Pipeline (24). The harmonized TCGA-BRCA data was downloaded on 7^th^ January 2020 from the Genomic Data Commons Data Portal using the R package ‘TCGAbiolinks’ (https://portal.gdc.cancer.gov/) (25). The end position of each isomiR feature is exclusive. Thus, the end position annotation was corrected by subtracting 1. For re-annotation of the data, an adaptation of miRBase version 22.1 was applied (26). All isoforms with the canonical 5’end of miR-193b-3p but regardless of their 3’end were summed up and considered as miR-193b-3p. Plate and tumour purity batch effects from sequencing were corrected with the ‘ComBat’ function of the R package ‘sva’ (https://rdrr.io/bioc/sva/man/ComBat.html) (27). Therefore, expression of miR-193b-3p for each patient was calculated as the sum of reads obtained from all 3’isomiRs with canonical seed sequence (28).

Normalized miRNA expression data of the METABRIC study (29,30) was obtained as arbitrary units from the EGA (https://www.ebi.ac.uk/ega/studies/EGAS00000000083, https://ega-archive.org/studies/EGAS00000000122). Data from both discovery and validation sets was merged into a single analysis of miRNA expression in patients. The data was array-based and did not allow discrimination between different microRNA isoforms.

For both datasets, PAM50 classification (31) was directly available in the clinical information, while TNBC status was defined as absence of ER, PR, and Her2 expression at the histological level. Only patients with available PAM50 as well as negative receptor status were considered, resulting in n=658 patients for TCGA and n=1293 patients for Metabric.

### Gene set enrichment analysis for TNBC patients

TCGA-BRCA cohort data after batch-correction and isomiR discrimination was used to investigate the correlation between miR-193b-3p expression and the activity of the selected pathways in patients. Batch-corrected expression of miR-193b was used as parameter and batch-corrected mRNA expression data was used as the input file. Spearman correlation coefficients between miR-193b and expressed genes were used as ranking metric and permutation was performed by phenotype. Default parameters of GSEA software (32,33) were applied.

## RESULTS

### miRNAs mildly regulate expression of proteins belonging to WNT/β-catenin, c-Met and Integrin signalling

In a previous study, we addressed miRNAs’ ability to regulate the cell cycle dependent on EGFR signalling (8). Here, we set out to identify miRNAs controlling metastatic traits in TNBC mediated by coordinated regulation of the c-Met, Integrin, and WNT/β-catenin signalling pathways. To investigate global effects, we employed a targeted proteomic approach, termed reverse phase protein array (RPPA). We started by selecting and classifying proteins belonging to these three pathways according to the KEGG Network database for c-Met (RTK signalling arm of map05200), integrin (map04510), and WNT (canonical WNT arm of map04310). Proteins originating from cognate mRNAs found expressed below 10 RPKM in an RNA sequencing dataset from MDA-MB-231 cells were excluded from further analyses. We then proceeded to perform antibody validation for specific detection of those proteins whose mRNA was expressed, identifying 62 antibodies appropriate for RPPA (supplementary table 1). We performed RPPA on a set of protein lysates derived from a gain-of-function assay where 800 miRNAs had been individually transfected in MDA-MB-231 cells (8). Subsequently, we used the limma test to quantify the impact of each miRNA on target proteins’ expression and compared to that to data obtained with miRNA mimic negative controls used in the same transfection batch. The global effect of miRNAs on all targets is summarized in a heatmap of log2 fold changes compared to the negative controls in Fig 1A. The limma test allowed us to maintain separate the information of the statistical significance and the amplitude of variation of target protein expression (Supplementary table 2 – all q-values and log2FC). All downstream analyses on the HTS data were performed only on miRNA-mediated effects having a q-value ≤0.001, thus maintaining a very stringent cutoff for statistical significance. Considering protein expression downregulation and upregulation separately, the average significant effects of miRNAs OE across all the proteins measured were -0.54 and +0.49 log2FC. In both directions we observed maximum effects reaching the absolute values of almost 3 log2FC (Fig 1B). In the HTS we identified a total of 5’435 significant interactions (out of 44’764). Of these, roughly 2/3 were downregulations and 1/3 upregulations (3804 and 1631, respectively). At the individual protein level, we noticed that the adaptor protein SHP-2, and the small GTPase p21-Rac were not significantly upregulated by any miRNA (Fig 1A, upper heatmap rugs, Supplementary fig 1). On the contrary, the remaining 60 targets were significantly upregulated and downregulated by at least one miRNA. However, these effects were not uniform, differing both in number of regulating miRNAs and their extent (Fig 1A, upper heatmap rugs, Supplementary fig 1). Indeed p53, Diap1, FAK1 showed greater log2FC variations compared to other targets, with averages above absolute value of 1 log2FC for both upregulations and downregulations (Supplementary fig 1). Focusing on the numbers of regulating miRNAs, low-density lipoprotein receptor-related protein 6 (LRP6) displayed a highly symmetrical regulation with 86 miRNAs upregulating its expression and 59 miRNAs repressing it. Another receptor protein, the EGF receptor (EGFR) was predominantly repressed by miRNAs OE (190 down vs 5 up). Conversely, the cell cycle negative regulator p27/Kip exhibited a predominant upregulation by multiple miRNAs (28 down vs 244 up) (Supplementary fig 2A). These strikingly different regulatory patterns prompted us to investigate further the presence of underlying biological features.

miRNA-mediated direct regulation of gene expression occurs predominantly via base-pairing with sequences located in the 3’ untranslated region (UTR) of target mRNAs. Thus, we wondered whether the differences in number of miRNAs regulating a target correlate with the length of the mRNA 3’ UTRs. Specifically, we reasoned that longer 3’UTRs would have a higher chance to harbour miRNA binding sites, thus negatively regulating the expression of the cognate protein. Meanwhile we expected that upregulation of protein expression upon miRNA OE would more likely be indirect and therefore unrelated to the length of mRNA 3’UTR. To address this, RNA-seq data from MDA-MB-231 cells was exploited to extract cell-specific 3’ UTR lengths, in nucleotides (nt.). There was no correlation between the number of miRNAs significantly regulating the target and the length of the 3’ UTR belonging to the mRNA. Employing an additional cutoff of |0.5| log2FC in defining a significant interaction, we found a positive trend between the number of miRNAs negatively regulating a target and the length of its 3’UTR (Supplementary fig. 2B, left panel). Unexpectedly, we observed instead a negative trend in correlation for the quantity of miRNAs upregulating protein expression (Supplementary fig. 2B, right panel). However, both these correlations were not statistically significant. Still, to validate if our RPPA results followed known patterns of protein expression regulation induced by miRNAs, we tested for predicted binding sites (BS) enrichment in the 3’ UTR of cognate RNAs (Agarwal 2015). Two independent algorithms were used to generate three separate datasets of predicted miRNA targets: TargetScan Conserved (TSC), TargetScan Non-conserved (TSNC), and microcosm (mC). The enrichment tests were performed independently on the list of miRNA-mediated repressions and on the one of the upregulations. We expected that the increase in protein expression upon miRNA OE would not be mediated by regulations via the 3’ UTR of target mRNAs and thus that there would be no significant enrichment in predicted binding sites. For the downregulating interactions, about half of the targets (32/62) presented a significant enrichment (p-value < 0.05) in at least one dataset and roughly one third of the targets (20/62) in two datasets. Among the upregulations, five targets showed a significant enrichment in one dataset while only one, MAPK14 (p38 protein), in two datasets (supplementary table 3).

These data show that the patterns of regulation that we uncovered followed known rules in miRNA-mediated gene expression control at the protein level. Importantly, we identified with high statistical confidence many mild interactions within the three pathways investigated, similarly to what was previously reported (5,6). Thus, we concluded that our RPPA dataset reliably quantified significant effects of miRNAs on proteins of interest. Next, we addressed miRNAs’ capability to regulate entire pathways, despite regulating only mildly individual targets.

### miRNAs coordinately control multiple pathways

The fine-tuning patterns identified in the RPPA HTS prompted us to further explore the function of miRNAs as regulators of pathways, shifting the focus from miRNA-protein to a miRNA-network context. In this new perspective, a two-step approach was taken: at first we reasoned that the biological effect of a miRNA on a pathway depends on the function that the regulated protein itself has. Then, once this information is integrated, the global effect of a miRNA on a pathway must be summed up based on its regulation of individual targets.

Therefore, for each target we assigned the pathway of origin as well as a putative positive or negative effect on said pathway based on literature and KEGG pathway descriptors (Fi g 1A lower rug and supplementary table 4). Then we defined that for each miRNA-protein pair, if the miRNA significantly downregulated the expression of a repressor of the pathway, the putative effect on the pathway would be positive. Conversely, if a miRNA downregulates an activator of the pathway, the effect would be negative. The opposite set of rules was applied for miRNAs upregulating their targets (Fig 1C). Focusing on the directionality of regulation, we defined as +1 a positive effect and -1 a negative effect on the pathway as mediated by the regulation of the probed target. Thus, following these rules, pathway-specific matrices of statistically significant log2FC, which we defined as miRNA-to-Protein (miR-P), were permuted into miRNA-to-Network (miR-N) matrices. These matrices represent the putative effect of a single miRNA on the pathway mediated by the individual probed target. To quantify the global effect of each miRNA on the complete pathway, we summed up the permuted values in the miR-N matrices and divided the obtained number by the quantity of targets probed within the pathway analysed. The newly defined Pathway Coregulatory (PC) score therefore describes the putative ability of a miRNA to regulate the whole network of proteins, normalized by the number of targets belonging to one pathway probed in the HTS (Fig 1C). The distribution of PC scores is biased by the number of proteins probed per pathway, as well as their putative role. Indeed, for WNT/β-catenin, where we probed 7 activators and 7 repressors, results show a symmetric distribution between repressing and activating miRNAs. On the contrary, for the other two pathways the imbalance caused by probing fewer repressors is shown in a distribution shifted toward repressive miRNA distributions (Fig 1D).

To identify high confidence miRNAs able to regulate the pathways, we tested the calculated PC scores against randomly generated ones. For that purpose, the miR-N matrix for each pathway was randomly permuted 10’000 times and, at each round, miRNA-specific random PC scores were generated. Then the actual PC score was tested against all randomly generated ones (Fig 1E). miRNAs whose actual PC score was significantly different compared to the randomly generated ones (alpha-level 5%) were regarded as actual pathway regulators (supplementary table 5). The effect of miRNAs was categorized into activators or repressors of the pathway if their actual PC score was significantly higher or lower than the random ones, respectively. The results of this analysis show that miRNAs regulating more than one pathway have either a consistent repressive or activating effect on all three pathways (Fig 1F). The only exceptions to this observed pattern are miR-409-3p, (repressing WNT, while activating integrin and c-Met) and miR-374b-3p (repressing WNT, activating integrin). For each separate pathway, two thirds of the miRNAs significantly negatively regulating it are coregulating in the same direction at least one other pathway, with 23 miRNAs regulating all three pathways. These results are similarly recapitulated for the positive regulations of c-Met and Integrin signalling pathways. Interestingly, despite displaying the most symmetric distribution of PC scores, the WNT pathway returned the lowest number of significantly upregulating miRNAs (Fig 1D and 1F).

Based on our PC score analysis, we concluded that miRNAs regulating one pathway have the tendency to concordantly regulate other interconnected pathways as well. In turn, this hinted at their ability to affect phenotypes even in the context of mild individual target regulation. Thus, we next sought to validate the capacity of these miRNAs to regulate phenotypes of TNBC by impinging on the selected pathways.

### miRNAs repress WNT/β-catenin and regulate proliferation upon pathway activation

We initially focused on validating miRNAs regulating the WNT/β-catenin signalling network where the pathway activity can be quantified based on transcriptional activity driven by TCF/LEF responsive elements. To test if miRNAs identified as repressors are indeed able to reduce the pathway activity, we performed gain-of-function experiment on MDA-MB-231 cells that harbour a stably integrated reporter (i.e., Firefly luciferase) under the control of TCF/LEF responsive elements. Here, six candidate miRNAs were tested: miR-103a-3p, miR-193b-3p, miR-409-3p, miR-494-3p, miR-889-3p, and miR-92b-3p. For all these miRNAs, the 3p arm of the miRNA precursor is the considered the guide miRNA according to miRBase (v22), and as such they will be further identified without the -3p suffix. As a positive control, we used iCRT-14 (inhibitor of Catenin Responsive Transcription-14), a small molecule that inhibits the interaction between the insulator proteins TCF/LEF and β-catenin, thus repressing transcription dependent on the latter (34). Luciferase activity was evaluated two days after transfection with miRNAs and 18hrs after stimulation with recombinant WNT3a. Compared to control conditions, miR-889 and miR-103a did not alter reporter gene activity. In contrast, miR-193b, -409, -494, and -92b significantly repressed the reporter gene activity by 50% or more (Fig 2A) indicating efficient suppression of the pathway. A similar level of repression of reporter gene activity was observed upon iCRT-14 treatment. This highlights the high accuracy of the PC score to predict miRNAs regulatory function on the pathway.

**Figure 2.**
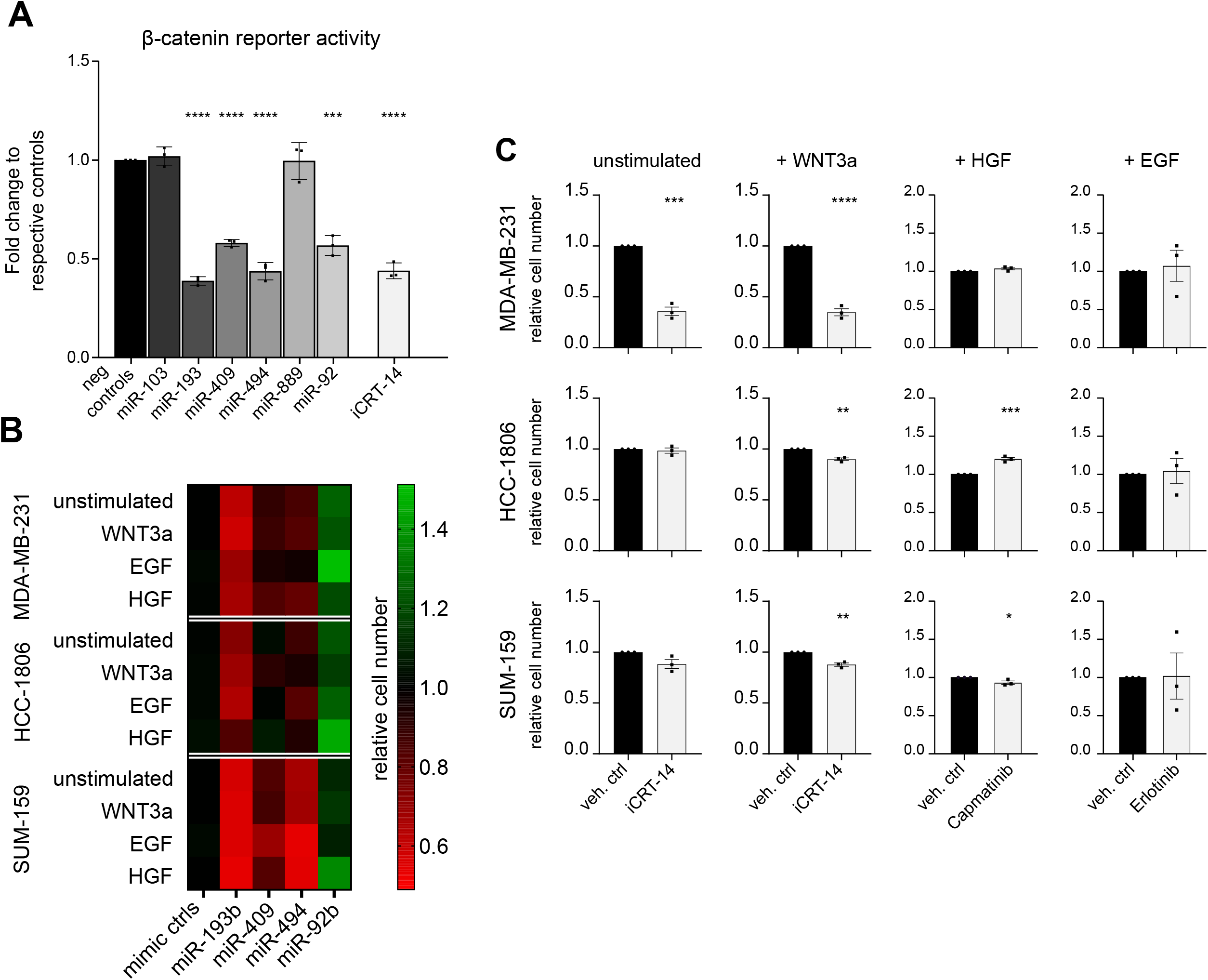
candidate miRNAs repress pathway activity and pathway-dependent growth. A. Stable isogenic Recombinant (SiR) MDA-MB-231 cells are transfected with miRNA mimics or treated with iCRT-14. 30 hrs later cells are stimulated with recombinant WNT3a. 18hrs later, FLuc and RLuc activities are assayed. The effect of miRNAs and iCRT-14 are shown on normalized luciferase activity relative to respective negative controls. Unpaired, two-tailed t-test on three independent SiR clones. P-values ***≤0.0001, * = 0.0168. B. MDA-MB-231, SUM-159, and HCC-1806 cells are transfected with miRNA mimics, five hours later medium is changed and, where marked, pathways are stimulated. 72hrs post-transfection cells growth is evaluated by nuclei counts. The effect of miRNAs is compared to two negative miRNA mimics. Each experiment is repeated in biological triplicate, with six technical replicates each. Growth reduction is represented in red and growth induction in green. Corresponding full bar charts are shown in Supplementary fig 3. C. MDA-MB-231, SUM-159, and HCC-1806 cells are treated with compounds or vehicle controls in combination with pathway stimulations, where marked. 72 hrs later cell growth is evaluated by nuclei counts. The effect of treatments is compared to relative vehicle (DMSO for iCRT-14 and Capmatinib, PBS for Erlotinib). Each experiment is repeated in biological triplicate, with six technical replicates each. Significance is calculated by unpaired, two-tailed t-test. P-values ****≤0.0001, ***≤0.001, **≤0.01, *≤0.05. non-significant are not marked.

One of the main phenotypes dependent on the activation of the canonical WNT pathway is cell cycle induction and cell proliferation. Thus, we asked whether the four miRNAs identified as repressors of the pathway could regulate cell growth accordingly. We investigated the effect of miRNAs and chemical inhibition of the pathway in two additional cell lines representing the same TNBC subtype (2). MDA-MB-231, SUM-159, and HCC-1806 cells were transfected with miRNA mimics and treated with recombinant WNT3a. Three days after transfection, their growth was compared to the one of controls by counting of nuclei as a proxy of proliferation. miR-193b consistently repressed proliferation of all three cell lines, both in WNT-stimulated and unstimulated conditions. miR-494 mildly but significantly reduced cell growth across all cell lines, while miR-409 displayed variable effects depending on the cell line and stimulation conditions. Unexpectedly, miR-92b upregulated proliferation (Fig 2B and Supplementary fig 3 for full bar charts of proliferation experiment). Chemical inhibition of the pathway with iCRT14 strongly repressed proliferation of MDA-MB-231 cells, both in stimulated and unstimulated conditions. However, in the other two cell lines, iCRT-14 treatment had no or little effect (Fig 2C and Supplementary fig 4). Considering the different effects caused by iCRT-14 across cell lines, we hypothesised that the three miRNAs could affect proliferation via different pathways in HCC-1806 and in SUM-159 cells. Hence, we evaluated the effect of these miRNAs on cell growth while stimulating c-Met and its co-interacting partner EGFR with recombinant HGF or EGF, respectively (Fig 2B). Additionally, we performed chemical inhibition of downstream signalling by treating cells with Capmatinib or Erlotinib (Fig 2C). The only miRNA retaining growth suppressive capabilities regardless of cell line or stimulation was miR-193b, except for HGF-treated HCC-1806 (Fig 2B). However, chemical inhibition of c-Met and EGFR signalling pathways did not cause similar effects in any cell line (Fig 2C and Supplementary fig 4).

Concluding, we validated the effect of three miRNAs as repressors of WNT/β-catenin signalling pathway, as well as their ability to suppress cell growth. One of the three miRNAs, miR-193b, displayed a strong phenotype which was not recapitulated by individual pathway suppression with chemical inhibitors. Therefore, we hypothesised that miR-193b functions by coordinate repression of the protein network and proceeded to investigate further phenotypes downstream of WNT/β-catenin and c-Met pathways.

### miR-193b induces apoptosis and represses migration and stem-like features of TNBC cell lines

To address miR-193b function in regulating apoptosis, we performed a gain-of-function experiment in MDA-MB-231 cells and assayed the percentage of apoptotic cells three days later. As hypothesised, miR-193b increased apoptosis significantly between 10- and 20-fold compared to negative controls. This was evident not only in unstimulated cells, but also when WNT and c-Met pathways were stimulated (Fig 3A). Conversely, none of the chemical inhibitors induced apoptosis to a similar extent (Fig 3B). Indeed, only Capmatinib treatment upon HGF stimulation caused a significant increase in apoptosis. However, the drug-effect was only minor, compared to the effect caused by miR-193b (Fig 3A and 3B).

**Figure 3.**
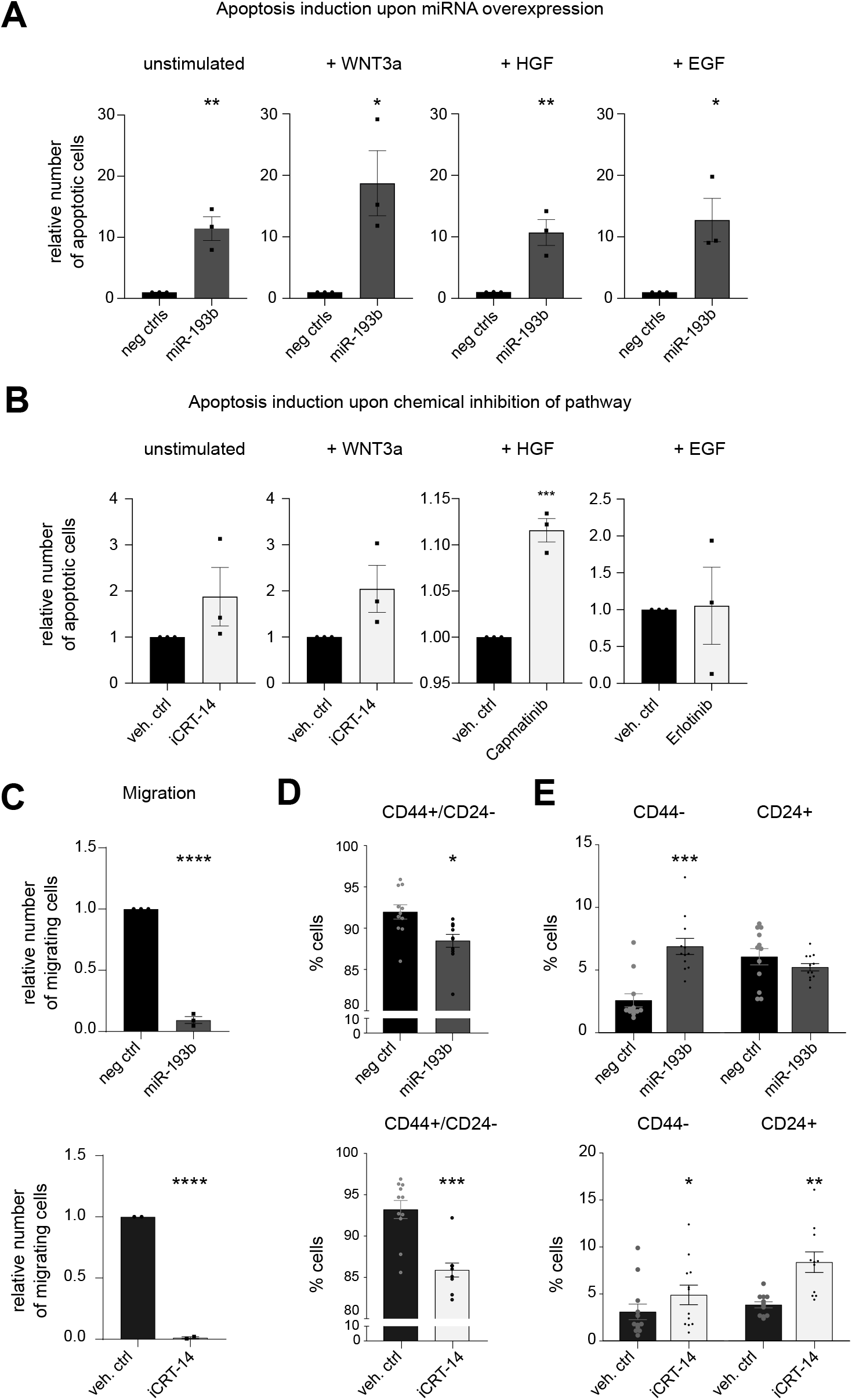
miR-193b regulates apoptosis, migration, and stemness in TNBC cell lines. **A** and **B** – MDA-MB-231, HCC-1806, and SUM-159 cells are treated and 72hrs later apoptosis is evaluated by Propodium Iodide positive nuclei over totals. For each condition, the effect of treatments is quantified relative to respective controls. Unstimulated conditions represent BSA-containing media to the same final concentration present in stimulation conditions, where it was used as carrier protein. **A)** The effect of miR-193b is compared to two negative miRNA mimics. **B)** The inhibitors in use are diluted in two different vehicles indicated in relevant graphs. Each experiment is repeated in biological triplicate, with six technical replicates each. Significance is calculated by unpaired, two-tailed t-test. P-values ***≤0.001, **≤0.01, *≤0.05. non-significant are not marked. C. Serum-starved MDA-MB-231 cells overexpressing miR-193b or treated with iCRT-14 are seeded in the upper compartment of a transwell system, with serum in the lower chamber as chemoattractant. 20 hrs later, migration relative to controls is evaluated by nuclei counts. The effect of miR-193b is tested against miRNA mimic negative control #2, and the effect of iCRT-14 against its vehicle, DMSO. The experiment is repeated in biological triplicate with three transwell parallel replicate each. Significance is calculated by unpaired, two-tailed t-test. P-values ****≤0.0001. **D** and **E** – FACS analysis of CD24 and CD44 surface marker expression in MDA-MB-231 cells 96 hrs post-transfection with miR-193b or iCRT-14 treatment, compared to respective controls. The experiment is repeated in six biological replicates, with two technical replicates each (all 12 datapoints plotted). Significance is calculated by paired, two-tailed t-test. P-values indicated on each graph. P-values ***≤0.001, **≤0.01, *≤0.05. non-significant are not marked. **D)** For each condition, the percentage of cells gated as CD44 positive and CD24 negative (stem-like population) is plotted. **E)** To highlight the differential effect of miR-193b and iCRT-14, the markers are displayed separate: for each condition, the percentage of cells gated as CD44 negative (left bars) or CD24 positive (right bars) are plotted.

The ability of miR-193b to reduce expression of multiple targets within the integrin pathway, including FAK, PAK, and Paxillin, hinted at a function for the miRNA to additionally affect cell motility. Thus, we next tested the effect of miR-193b on migration of MDA-MB-231. Serum-starved cells were transferred in the upper chamber of a transwell system 72 hrs after miRNA overexpression and were allowed to migrate for 20 hrs toward serum-containing medium. Migrated cells were quantified by nuclei count and normalised for seeding differences. miR-193b nearly abolished migration toward serum, to a similar extent as cells treated with iCRT-14 (Fig 3C).

Importantly, WNT pathway is strongly associated with maintenance of stemness in diverse cellular contexts, such as embryonic stem cells, intestinal adult cells, and breast cancer (35,36). The stem-like population of cells in breast cancer is characterised by high expression of CD44 and low or no expression of CD24 (CD44+/CD24-). Thus, we tested by Fluorescence Activated Cell Sorting (FACS) the expression of these two surface markers four days post-transfection of miR-193b or chemical inhibition of the pathway. Both treatments significantly reduced the stem-like population (CD44+/CD24-) (Fig 3D). However, analysis of the individual markers showed that miR-193b affected predominantly the expression of CD44, significantly increasing the population of CD44- cells compared to a miRNA mimic control. Oppositely, iCRT-14 treatment did not significantly affect the CD44 expressing population, rather increased the CD24+ cell population (Fig 3E).

Hence, we demonstrated that miR-193b overexpression *in vitro* limits not only proliferation, but also additional phenotypes linked with TNBC metastatic traits. Importantly, some of these phenotypes were not recapitulated by individual pathway inhibition, indicating how miR-193b coordinately regulates multiple signalling pathways collectively driving aggressive cancer phenotypes. To further validate the importance of miR-193b in the context of TNBC, we next analysed miRNA expression data derived from BC patients.

### In TNBC patients, miR-193b has lower expression and regulates WNT/β-catenin and c-Met signalling pathways

To address miR-193b regulatory function on these pathways and thus the aggressiveness of the disease, we analysed miR-193b expression in two independent datasets of breast cancer patients (TCGA BRCA and METABRIC) (24,29,30) stratifying them by histological status or by PAM50 classifier. In both datasets miR-193b expression was significantly lower in patients of TNBC subtype (Fig 4A). Similarly, miR-193b was significantly less expressed in patients classified as Basal by PAM50 signature compared to the other three subtypes (Fig 4B).

**Figure 4.**
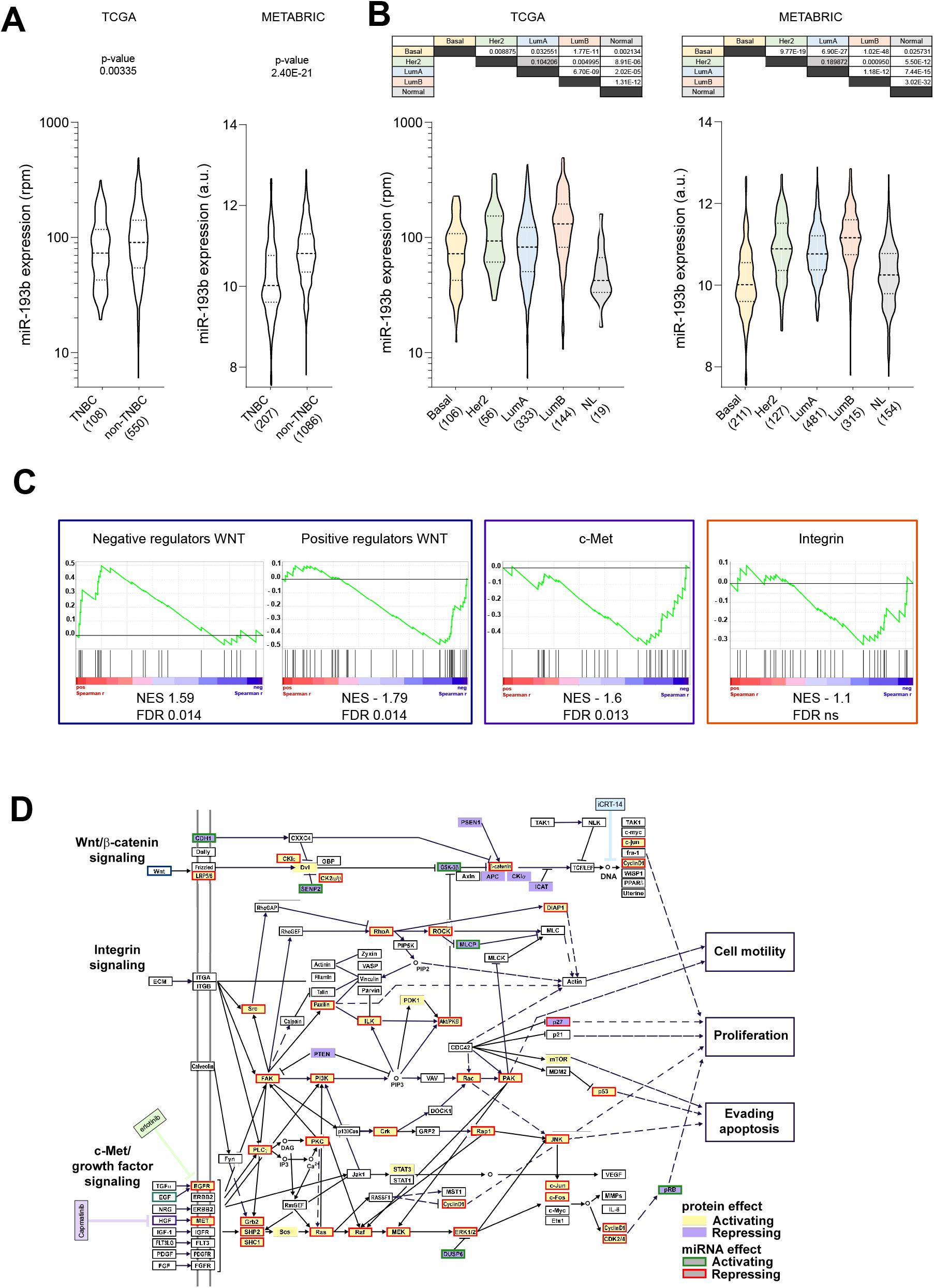
miR-193b expression in BRCA patients is associated with aggressiveness and gene sets of signalling pathways of interest. **A** and **B** – Violin plots of miR-193b expression in two BRCA datasets. TCGA BRCA dataset quantification is in reads per million (rpm). Metabric dataset quantification is in arbitrary units (a.u.) Dashed and dotted lines within violins represent the median and quartiles of the distributions. A. miR-193b expression stratifying patients by receptor expression status in TCGA (left) and METABRIC (right) datasets. Within each dataset, number of patients belonging to the TNBC or non-TNBC classification are written in parentheses at the x-axes. Exact p-values are indicated. B. miR-193b expression in two BRCA datasets stratifying patients by PAM50 classification into Basal, Her2, Luminal A (LumA), and Luminal B (LumB). Reciprocal exact p-values are indicated in the table above graph. Except the ones marked in grey, all the others are below ≤0.05. C. Gene Set Enrichment Analysis of gene lists for positive and negative regulators of WNT signalling (blue box), c-Met signalling (purple box), and Integrin signalling (orange box). Normalized enrichment scores (NES) and statistical significance by false discovery rates (FDR) are indicated below every signature. D. Effect of miR-193b on the three signalling pathways, integrated according to their KEGG maps with downstream phenotypes. Target proteins probed in the HTS are shaded in lilac or pale yellow when they are repressors or activators of the pathways, respectively. miR-193b repressive or activating effect on the pathways is represented by a box around the proteins significantly regulated, of red or green colour, respectively. The chemical inhibitors’ activities are highlighted in blue (iCRT-14), purple (Capmatinib), and green (Erlotinib).

To consolidate the functional role of miR-193b as repressor of the pathways of interest, we performed a gene set enrichment analysis (GSEA) on patients’ gene expression data from the TCGA BRCA dataset. First, we selected TNBC patients by histological status. Then, we correlated miR-193b expression with that of all genes expressed, ranking them from highest to lowest by spearman correlation coefficient. Then, we compiled four lists of genes based on Biocarta (c-Met and Integrin signalling), and KEGG (positive and negative regulators of WNT/β-catenin signalling). GSEA indicated that genes encoding for negative regulators of WNT/β-catenin were enriched among the positively correlating genes (Fig 4C, first plot). Conversely, signatures of positive regulators of WNT (Fig 4C, second plot) and c-Met signalling (Fig 4C, third plot) were enriched among the negatively correlated genes. The same trend was observed for the Integrin signalling pathway, albeit not reaching significance (Fig 4C, fourth plot). Hence, these results further support miR-193b role as a regulator of the investigated pathways in TNBC patients.

In conclusion, analysing two independent datasets we have shown that miR-193b expression is the lowest in patients diagnosed with the most aggressive subtypes of breast cancer. Additionally, we confirmed *in vivo* the function of miR-193b as a negative regulator of WNT/β-catenin and c-Met signalling pathways. Therefore, we concluded that miR-193b is a master regulator of these pathways associated with metastatic traits in TNBC.

## DISCUSSION

In this project we aimed to identify miRNAs able to regulate c-Met, Integrin, and WNT/β-catenin signalling pathways, thus coordinately controlling phenotypes associated with aggressive traits in TNBC, such as growth, migration, and stem-like features. The effect of miRNAs on the proteins belonging to these three pathways was analysed using a targeted proteomic approach, RPPA. Selecting a very stringent statistical threshold, miRNAs were scored for their putative effect on the selected pathways. The validity of the Pathway Coregulatory score was demonstrated by a marked downregulation of WNT pathway activity by four out of six miRNAs tested with negative PC scores for WNT. Then, we characterised one of the top-scoring miRNAs, miR-193b, demonstrating its ability to regulate the signalling pathways in TNBC patients’ datasets and *in vitro* phenotypes dependent on their activity.

The proteomics approach we employed addresses the function of miRNAs at the protein rather than mRNA expression level. In the context of ER+ breast cancer, mass spectrometry (MS) coupled with iTRAQ (isobaric tag for relative and absolute quantification) has previously been exploited to identify in an unbiased fashion, the targets of miR-193b at the protein level (37). Gene expression analysis upon miR-193b overexpression showed that only a minority (13%) of the proteins identified had a matched repression of its mRNA molecule (37). This highlights the importance of considering proteins as they are the functional effectors of signalling pathways and their respectively regulated phenotypes. To overcome the limitations of MS in throughput regarding the number of miRNAs to be investigated, we employed RPPA in this study. We undertook a bootstrap analysis pipeline to quantify effects on pathways and proved that the significant co-regulatory effects were greater than random ones. This approach allowed us to cope with potential biases of this targeted proteomic approach. While this does not exclude the possibility that the calculated PC scores are still biased due to the selection of probed proteins, the benchmarking analysis of negative regulators of the WNT pathway demonstrates the validity of our setup. Our analysis of multiple pathways showed that miRNAs negatively regulating one tended to have the same function also on neighbouring pathways, thus supporting the concept that miRNAs can effectively regulate complex phenotypes even if their effects are rather mild at the single-target level.

The role of the miRNome on the WNT/β-catenin signalling pathway had been previously studied by Anton and colleagues upon transfecting individual miRNA in HEK293T cells together with a TOP/Flash reporter (38). The list of miRNAs repressing WNT/β-catenin signalling in our screen were thus compared to those found in the published screen: four miRNAs were concordantly present in the two lists - miR-193b, miR-409, and miR-28 were all described as mild repressors of the reporter gene. On the other hand, miR-181d was characterized as activator. All four have instead been characterised as negative regulators in our analysis. Experimentally, in MDA-MB-231, miR-193b and miR-409 downregulated the signalling pathway, whilst neither miR-28 nor miR-181d could regulate reporter gene expression (unpublished data). Some experimental differences could partially explain the results: e.g. Anton and colleagues transfected miRNA mimics at 40nM, possibly rendering the screening more prone to off-target and indirect effects when compared to the 25nM concentration used in our RPPA screening. Nevertheless, biological differences could also explain these divergent results: within a cell system, the presence or absence of specific transcripts, as well as their abundance can deeply affect the role of a miRNA. Therefore, while supporting some of our findings, the screening from Anton and colleagues emphasizes that the effect of miRNAs should be considered in the context of cell and tissue types.

The capacity of miR-193b to regulate concordantly several pathways is of high importance, particularly when compared to the effects we observed with chemical inhibitors of the individual pathways, showing that by targeting at multiple levels, a miRNA exerts a stronger functional output than an upstream inhibition. The ability of miR-193b to target different pathways and thus coordinately regulate a phenotype is exemplified by the fact that the other two tested cell lines (HCC-1806 and SUM-159) do not respond to inhibition of WNT pathway to the same extent as MDA-MB-231. Thus, their proliferation probably is not dependent on this pathway. Nevertheless, miR-193b strongly repressed proliferation also in these cell lines. Interestingly, the effect of miR-193b was greatly reduced in HCC-1806 when cells were concomitantly stimulated with HGF. This is a trend similar to how their proliferation slightly increases when treated with Campatinib, hinting possibly at a specific c-Met-EGFR crosstalk in this cell line.

Our key finding is the capacity of miR-193b to regulate WNT/β-catenin and c-Met pathway in TNBC, both *in vitro* and in gene expression signatures derived from patients. Previous findings had circumscribed the role of miR-193b as a repressor of individual targets in TNBC, such as its ability to individually downregulate urokinase-type plasminogen activator (uPA) (39), or dimethylarginine dimethylaminohydrolase 1 (DDAH1) (40). However, possibly pursuing a more physiological avenue, we present the concept that miR-193b exerts its function as tumour suppressor by coordinately regulating entire pathways that are relevant for the acquisition and maintenance of aggressive features. Supported by literature, we recapitulate its effect on growth and migration previously identified (39,40), and we further characterised its function on apoptosis induction and repression of stem-cell like features such as the expression of CD24 and CD44 surface markers. Chemical inhibition of WNT pathway via iCRT-14 treatment mimics some of these phenotypes, supporting the idea that they are partially downstream WNT pathway specifically. However, based on the discrepancies seen in the response to miRNAs and to chemical inhibition of WNT/β-catenin signalling in the three cell lines assayed, we hypothesize: a) that miRNAs regulate proliferation by affecting multiple pathways, and b) that the proliferation of different cell lines, also riddled by their particular mutation statuses, might depend on different pathways. We speculate that the same principles apply also *in vivo*, where intratumoral heterogeneity could be promoted by loss of miR-193b.

The WNT/β-catenin signalling pathway has recently gained attention for its effects on TNBC, despite absence of recurrent β-catenin mutations or classical genetic lesions associated to this pathway’s overactivation, such as APC loss in colorectal cancer (41). Additionally, the pathway was shown to be activated in basal-like breast cancers (akin to TNBC) where it was associated with worse prognosis (42). In another study, TNBC patients with a WNT-dependent gene expression signature presented higher rates of lung and brain metastases (43). At present, the causative role for enhanced activation of WNT/β-catenin signalling has not yet been pinned down to either a common mutation or genomic alteration. It is thus tempting to speculate that its regulation could in part be mediated by miR-193b that we, and others, have found expressed at lower levels in TNBC in patients’ derived gene expression profiles. Additionally, the ability of miR-193b to repress multiple pathways and its reduced expression in TNBC could explain those findings that show how TNBC outcome depends on a combination of deregulated pathways, such as Wnt/β-catenin, c-Met, and CXCL12/CXCR4 (44).

## CONCLUSIONS

In summary, we developed a new network analysis to unravel miRNAs functional relevance on signalling pathways regulating metastatic traits in TNBC. Focusing on WNT/β-catenin, c-Met, and Integrin pathways, we identified 23 miRNAs able to repress them in a coordinated fashion. We broadly validated across TNBC cell lines the phenotypic effects of our top candidate, miR-193b-3p. We demonstrated that miR-193b affects phenotypes differently than chemical inhibitors of individual pathways, proving its ability to target them at multiple levels. Ultimately, we showed how TNBC and basal patients display the lowest miR-193b expression, thus highlighting a potential mechanism that this tumour type employs to activate pro-metastatic signalling pathways.

## FIGURE LEGENDS

**Supplementary figure 1.**
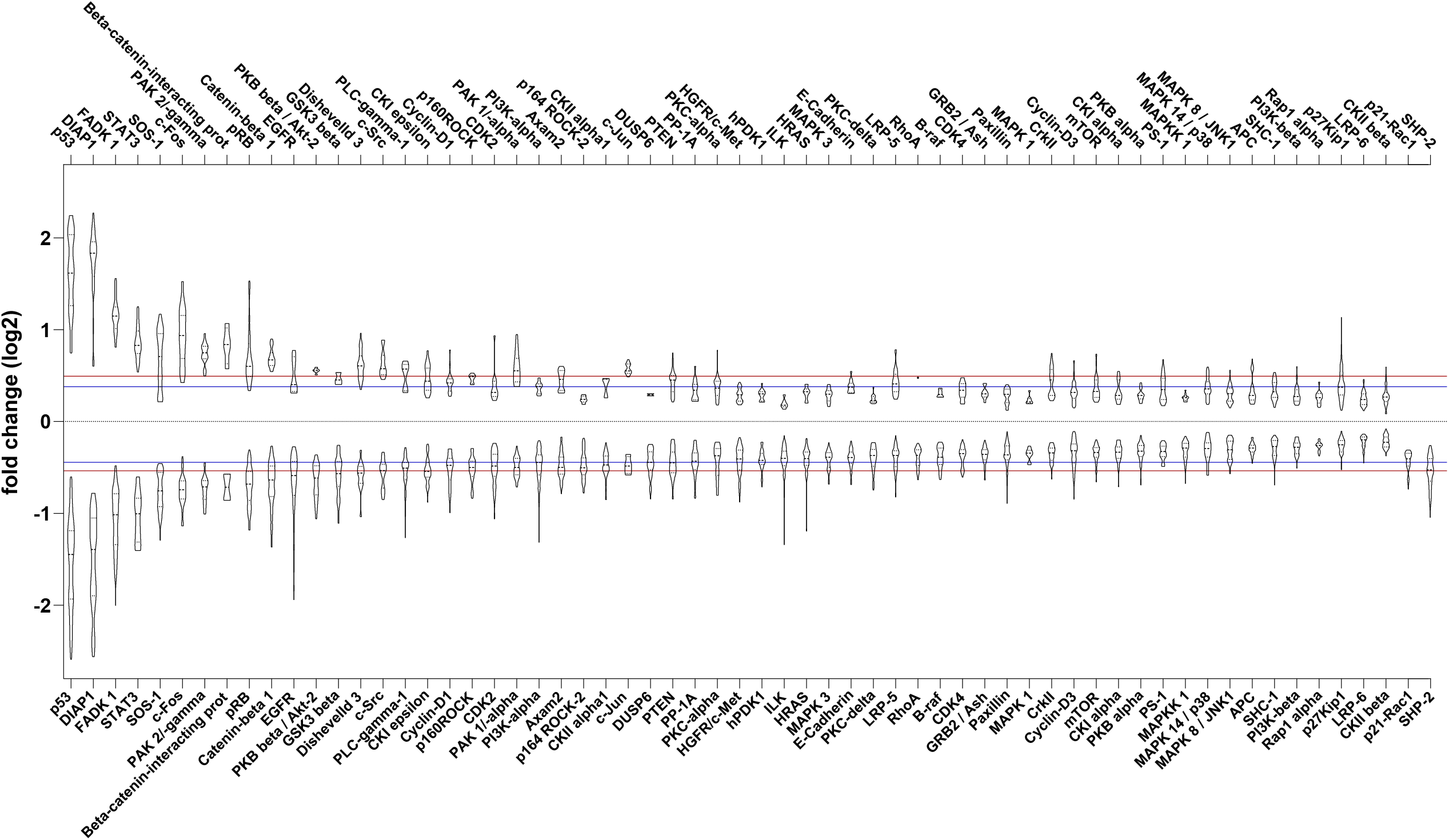
Quantifying effects of miRNAs library overexpression on individual targets. Violin plots of all significant (q ≤0.001) positive or negative log2FCs computed by limma testing, for each individual protein assayed. Proteins are ranked by average negative log2FC. p21-Rac, and SHP-2 are displayed at the end of the distribution since they are not significantly upregulated by any miRNA. Within each violin, dashed lines represent the medians and dotted lines represent the quartiles of the distributions. Horizontal red and blue lines represent the averages and medians, respectively, computed from the whole HTS.

**Supplementary figure 2.**
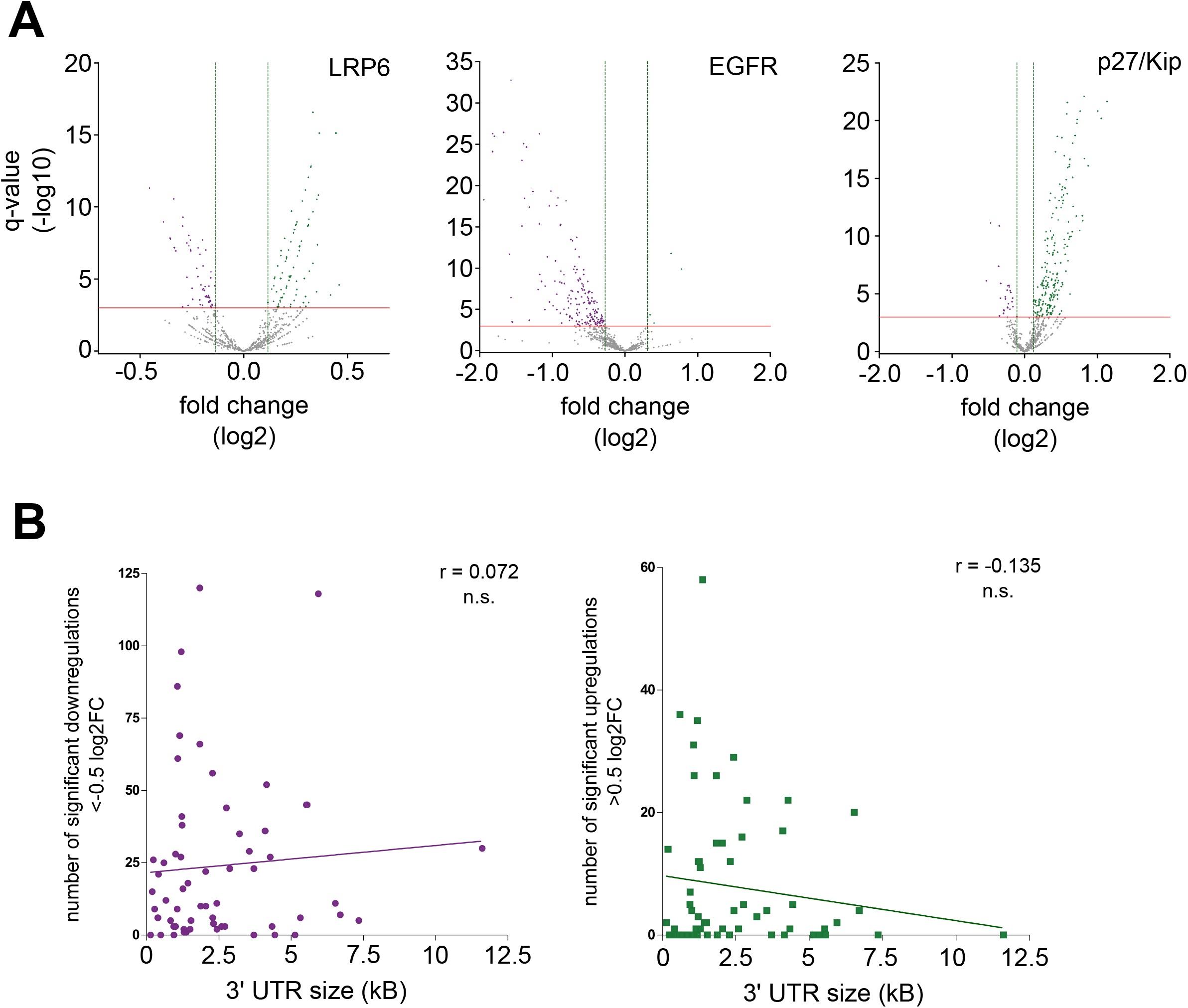
Addressing biological features of transcript regulating protein expression patterns. A. Effect of miRNAs on selected targets is represented in three volcano plots. The fold change of protein expression (in log2) compared to control miRNA mimics is plotted on the x-axes, q-values (in log10) are plotted on the y-axes. Red horizontal lines identify the q-value cutoff used for the downstream analyses (q ≤0.001), vertical dotted lines represent for each target the minimum log2FC at which the cutoff was passed. Statistically significant interactions are in purple (downregulations) or green (upregulations). The three targets displayed are Low-density lipoprotein receptor-related protein 6 (LRP6) (left panel), EGF receptor (EGFR) (middle), and p27/Kip (right). B. Effect of mRNA 3’UTR sizes on miRNA-mediated regulation. For each target assayed, the length of the 3’UTR of the cognate mRNAs is extracted from MDA-MB-231 sequencing data. Sizes of 3’UTRs on the x-axes (in nucleotides – nt.) are then plotted against the number of miRNAs significantly downregulating (left panel) or upregulating (right panel) the expression of the target proteins by at least an absolute value of 0.5 log2FC. For each distribution, Pearson r is computed and in both cases the p-value does not indicate a significant correlation.

**Supplementary figure 3.**
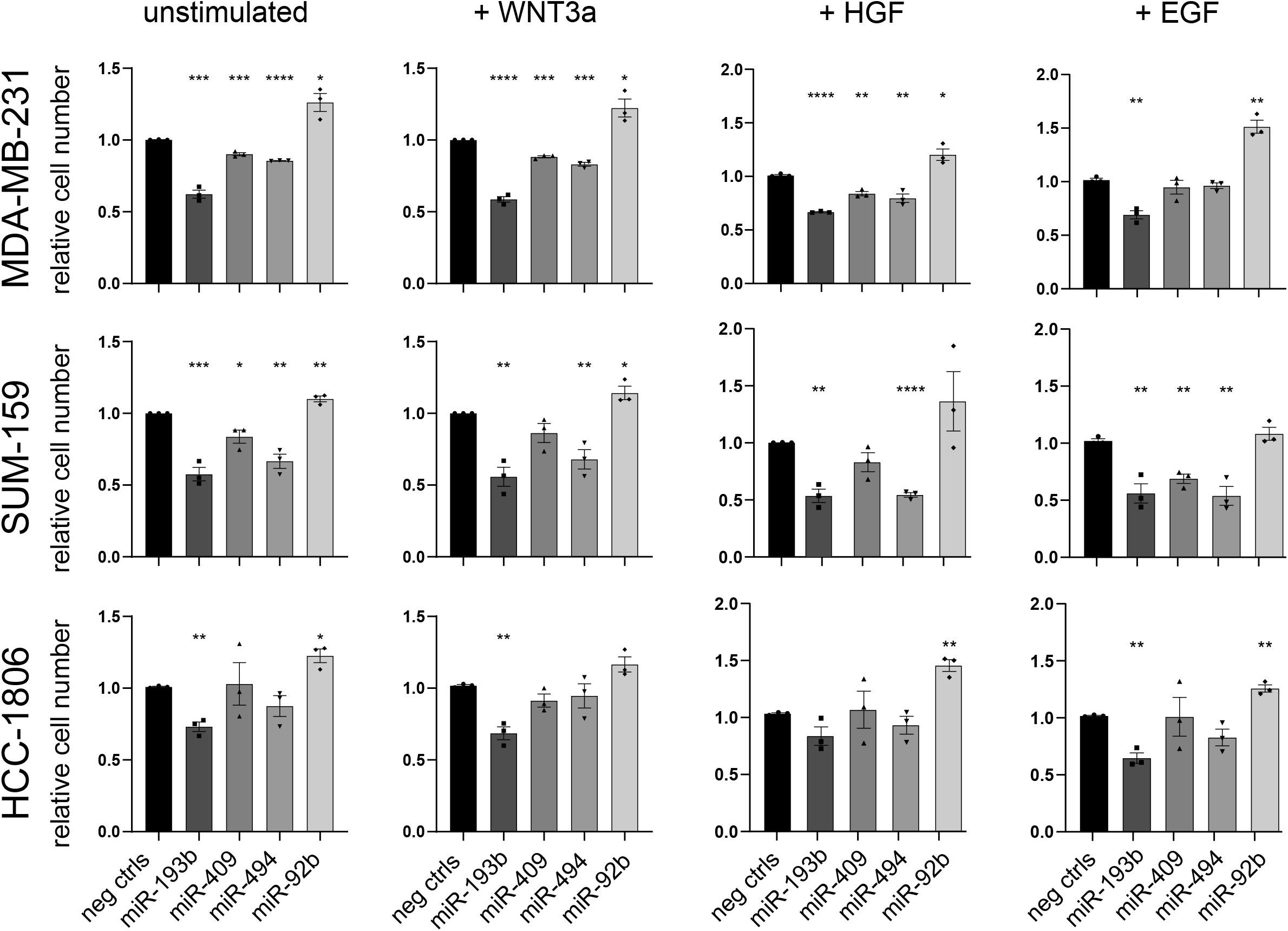
candidate miRNAs effect on cell growth. Data represented in main figure 2B is represented here as a bar chart to allow for individual value and statistically significance inspection. MDA-MB-231, SUM-159, and HCC-1806 cells are transfected with miRNA mimics, five hours later medium is changed and, where marked, pathways are stimulated. 72hrs post-transfection cells growth is evaluated by nuclei counts. The effect of miRNAs is compared to two negative miRNA mimics. Each experiment is repeated in biological triplicate, with six technical replicates each. Significance is calculated by unpaired, two-tailed t-test. P-values ****≤0.0001, ***≤0.001, **≤0.01, *≤0.05. non-significant are not marked.

**Supplementary figure 4.**
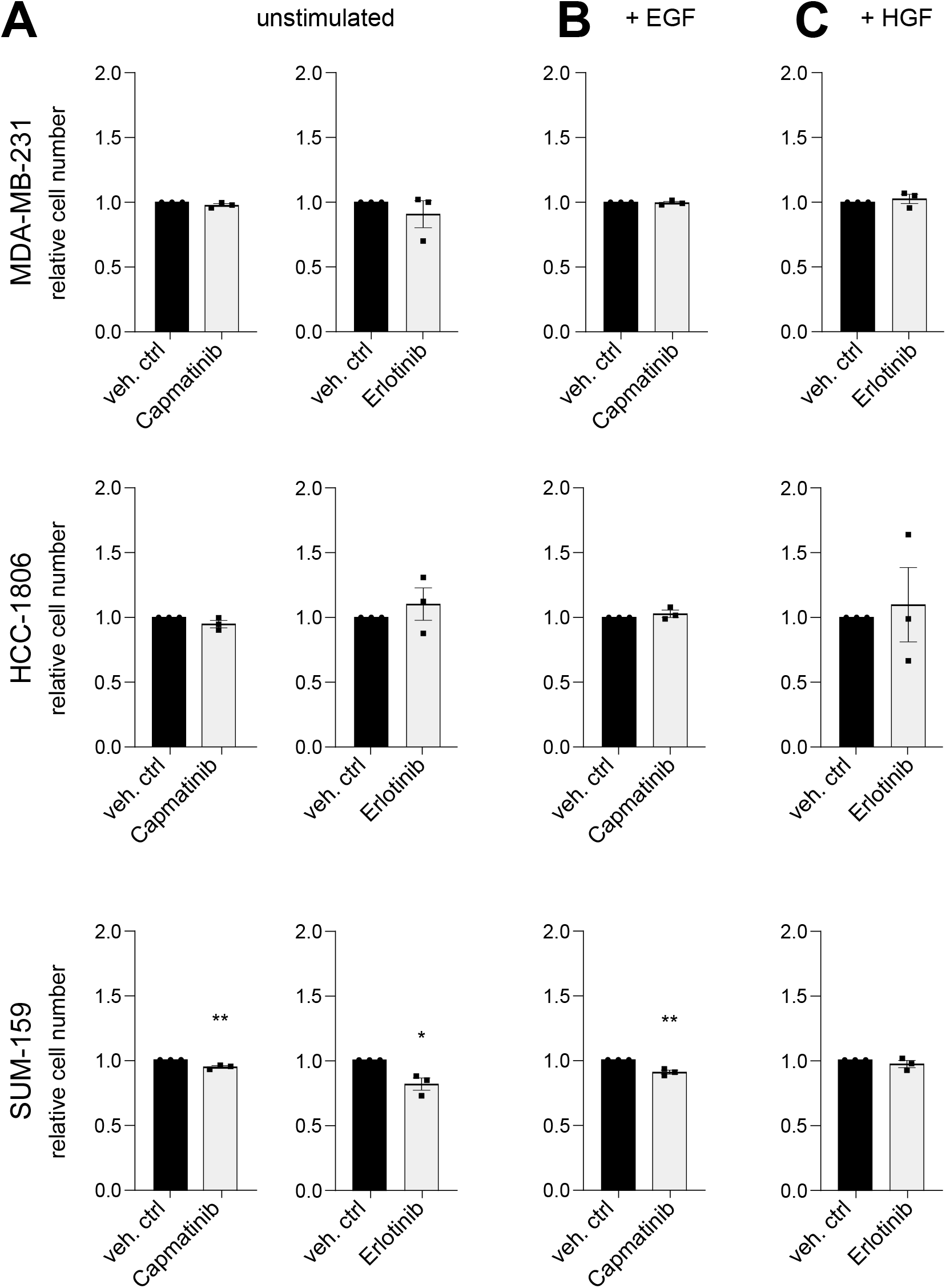
chemical inhibition of the pathway with reciprocal stimulations. **A** – **C** – MDA-MB-231, SUM-159, and HCC-1806 cells are treated with compounds or vehicle controls in combination with pathway stimulations, where marked. 72hrs later cells growth is evaluated by nuclei counts. For each condition, the effect of treatments is quantified relative to respective controls. The inhibitors in use are diluted in two different vehicles indicated in relevant graphs. The effect of treatments is compared to relative vehicle (DMSO for Capmatinib, PBS for Erlotinib). Each experiment is repeated in biological triplicate, with six technical replicates each. Significance is calculated by unpaired, two-tailed t-test. P-values **≤0.01, *≤0.05. non-significant are not marked.

## ACKNOWLEDGMENTS

We thank Angelika Wörner, Corinna Becki, and Daniela Heiss (Molecular Genome Analysis, DKFZ) for excellent technical support, Rita Schatten and Birgit Kaiser (Genomics and Proteomics Core faciities, DKFZ) for experimental services, and the Microscopy and FACS core facilities for providing instruments and technical assistance. We thank Sven Diederichs (DKFZ, Heidelberg) for discussions and Edward W. Roberts (CRUK Beatson Institute) for constructive feedback during manuscript preparation. Funding: CG was supported by a doctoral fellowship of the German–Israeli Helmholtz Research School in Cancer Biology. Additionally, this work was supported by the Landesstiftung Baden-Württemberg (BWST_NCRNA_035) and the German Federal Ministry of Education and Research (e:Med FKZ:031A429).

## AUTHOR CONTRIBUTIONS (<u>CRediT statement</u>)

CG Conceptualization, Methodology, Investigation, Formal Analysis, Data Curation, Visualization, Writing - Original Draft, Writing - Review & Editing, Project Administration

JJ Validation, Investigation, Formal Analysis

AW Software, Formal Analysis

SI Software, Formal Analysis

RW Resources, Methodology

SU Resources

HM Resources

OS Resources, Conceptualization

YY Supervision, Funding acquisition

TB Supervision

UK Supervision, Methodology

CK Conceptualization, Supervision, Formal Analysis, Visualization, Writing - Review & Editing

SW Conceptualization, Supervision, Funding acquisition, Writing - Review & Editing

