## Supplementary File 1 - code for "Coordinated regulation of WNT/β-catenin, c-Met, and Integrin signalling pathways by miR-193b controls triple negative breast cancer metastatic traits": 20210510_Supplementary_File_1_Code.html

PC score calculation


### PC score calculation

###### *December 16, 2017*

- 1. Read in data
  - 1.1 Read in data matrices from differential expression testing
  - 1.2 Read in collected literature knowledge of protein effect on WNT, MET and Integrin pathways
- 2. Protein regulation matrices
  - 2.1 Protein regulation matrix
  - 2.2 Pathway-specific protein regulation matrices
- 3. Putative effects on pathways
- 4. Pathway coregulation score
  - 4.1 Calculation of PC score
  - 4.2 Permutation testing
- 5. Session Info

#### 1. Read in data

##### 1.1 Read in data matrices from differential expression testing

Multiple testing adjusted p-values from differential expression test were filtered to those 722 miRNAs showing an adjusted p-value < 0.001 for at least one protein. Fold changes are used to consider up/downregulation individually.

```
dat = read.csv(paste(dir, "q_val_matrix_final0.001_ordered.csv", sep = ""), 
               row.names = 1,
               stringsAsFactors = FALSE)
fc = read.csv(paste(dir, "Mdiff_all_arrays_final0.001_ordered.csv", sep = ""), 
               row.names = 1,
               stringsAsFactors = FALSE)
```

##### 1.2 Read in collected literature knowledge of protein effect on WNT, MET and Integrin pathways

Putative protein effects on WNT, MET and Integrin pathways were collected from literature and summarized as positive or negative effects, respectively.

```
PW_effects = read.csv(paste(dir, "PW_prot_effects.csv", sep = ""), 
                      sep = "\t",
                      stringsAsFactors = FALSE)

prot_effect_WNT = PW_effects[-1,c(1,3)][1:14,]
prot_effect_MET = PW_effects[-1,c(4,6)][1:30,]
prot_effect_Int = PW_effects[-1,c(7,9)][1:34,]

#rename MTOR/FRAP1
prot_effect_MET[grep("MTOR", prot_effect_MET[,1]),1] = "FRAP1"
```

#### 2. Protein regulation matrices

##### 2.1 Protein regulation matrix

miRNA regulation is considered significant based on a multiple testing adjusted p-value < 0.001. Regulation matrix is discretized based on significant up/downregulation.

```
#regulation matrix
reg_mat = matrix(data = 0, ncol = ncol(dat), nrow = nrow(dat))
rownames(reg_mat) = rownames(dat)
colnames(reg_mat) = colnames(dat)
reg_mat[dat < 0.001 & fc < 0] = -1
reg_mat[dat < 0.001 & fc > 0] = 1
```

##### 2.2 Pathway-specific protein regulation matrices

Discretized regulation matrices are generated for WNT, MET and Integrin pathways.

```
#pathway regulation matrices
pw_reg <- function(prot_effect, reg_matrix)
{
    ind_reg = vector()
    for(j in 1: dim(prot_effect)[1])
    {ind_reg[j] = grep(prot_effect[j,1], colnames(reg_matrix))}
    reg_mat_pw = reg_matrix[,ind_reg] 
    return(reg_mat_pw)
}
reg_mat_WNT = pw_reg(prot_effect_WNT, reg_mat)
reg_mat_MET = pw_reg(prot_effect_MET, reg_mat)
reg_mat_Integrin = pw_reg(prot_effect_Int, reg_mat)
```

#### 3. Putative effects on pathways

Putative effects of miRNAs on pathways are summarized by combination of measured miRNA-protein effects and literature derived protein-pathway effects.

```
put_pw_effect <- function(reg_mat_pw, prot_effect)
{
  reg_mat_comb = reg_mat_pw
    for(s in 1:dim(prot_effect)[1])
    {  for(t in 1: dim(reg_mat_pw)[1])
        {if(reg_mat_pw[t,s] == 0)
            {reg_mat_comb[t,s] = 0 
        }else if(reg_mat_pw[t,s] == -1 & prot_effect[s,2] == "pos"){
            reg_mat_comb[t,s] = -1  
        }else if(reg_mat_pw[t,s] == 1 & prot_effect[s,2] == "pos"){
            reg_mat_comb[t,s] = 1  
        }else if(reg_mat_pw[t,s] == -1 & prot_effect[s,2] == "neg"){
            reg_mat_comb[t,s] = 1  
        }else if(reg_mat_pw[t,s] == 1 & prot_effect[s,2] == "neg"){
            reg_mat_comb[t,s] = -1  
        }
        }
    } 
  return(reg_mat_comb)
}
reg_mat_WNT_comb = put_pw_effect(reg_mat_WNT, prot_effect_WNT)
reg_mat_MET_comb = put_pw_effect(reg_mat_MET, prot_effect_MET)
reg_mat_Integrin_comb = put_pw_effect(reg_mat_Integrin, prot_effect_Int)
```

#### 4. Pathway coregulation score

A pathway coregulation (PC) score is defined for each miRNA as the sum of all measured miRNA-mediated effects on the pathway weighted by the number of measured proteins in the pathway.

##### 4.1 Calculation of PC score

The PC score is calculated for WNT, MET and Integrin pathways.

```
PC_score <- function(reg_mat_comb)
{
   PC_score = vector()
   for(s in 1: dim(reg_mat_comb)[1])
   {PC_score[s] = sum(reg_mat_comb[s,])/dim(reg_mat_comb)[2]}
   names(PC_score) = rownames(reg_mat_comb)
   return(PC_score)
}
PC_WNT = PC_score(reg_mat_WNT_comb)
PC_MET = PC_score(reg_mat_MET_comb)
PC_Int = PC_score(reg_mat_Integrin_comb)
```

##### 4.2 Permutation testing

Permutation testing is performed to assess the miRNA-wise probability distribution of PC scores by 10000x resampling the putative miRNA-pathway interactions for each protein. PC scores are considered significant based on an alpha-level of 5%, in addition the absolute PC score is assessed as distance from the mean of the permuted PC scores.

```
#permutation parameters
set.seed(308) 
n_perm = 10000

#permutation matrix
perm_mat <- function(reg_mat_comb, n_perm)
{
    PC_perm_mat = matrix(ncol = dim(reg_mat_comb)[1], nrow = n_perm)
    colnames(PC_perm_mat) = rownames(reg_mat_comb)
    rownames(PC_perm_mat) = paste("perm run", 1:n_perm, sep = " ")
    for(k in 1: n_perm)
    {   perm_col_mat = matrix(ncol = dim(reg_mat_comb)[2], nrow = dim(reg_mat_comb)[1])
        for(i in 1: dim(perm_col_mat)[2])
        {perm_col_mat[,i] = sample(reg_mat_comb[,i], size = dim(reg_mat_comb)[1], replace = FALSE)}
        PC_perm_mat[k,] = PC_score(perm_col_mat)
    }
    return(PC_perm_mat)
}
PC_perm_mat_WNT = perm_mat(reg_mat_WNT_comb, n_perm)
PC_perm_mat_MET = perm_mat(reg_mat_MET_comb, n_perm)
PC_perm_mat_Int = perm_mat(reg_mat_Integrin_comb, n_perm)

#thresholding
sign_PC <- function(PC_perm_mat, PC)
{
    vec_sign = vector()
    dval = vector() 
    PC_perm_mat_st = PC_perm_mat
    for(j in 1: dim(PC_perm_mat)[2])
    {
        PC_perm_mat_st[,j] = (PC_perm_mat[,j] - mean(PC_perm_mat[,j]))
        if((PC[j] - mean(PC_perm_mat[,j]) <= quantile(PC_perm_mat_st[,j], c(0.025))))
            {   vec_sign[j] = -1  
                dval[j] = PC[j] - mean(PC_perm_mat[,j])
            }else if(PC[j] - mean(PC_perm_mat[,j]) > quantile(PC_perm_mat_st[,j], c(0.975))){ 
                vec_sign[j] = 1 
                dval[j] = PC[j] - mean(PC_perm_mat[,j])
            }else{
                vec_sign[j] = 0    
                dval[j] = PC[j] - mean(PC_perm_mat[,j])
            }
    }
    vec_sign = cbind(vec_sign, dval)
    rownames(vec_sign) = colnames(PC_perm_mat)
    vec_sign = vec_sign[order(vec_sign[,1], vec_sign[,2]),]
    colnames(vec_sign) = c("sign", "dist")
    return(vec_sign)
}
WNT_sign_PC = sign_PC(PC_perm_mat_WNT, PC_WNT)
MET_sign_PC = sign_PC(PC_perm_mat_MET, PC_MET)
Integrin_sign_PC = sign_PC(PC_perm_mat_Int, PC_Int)
```

#### 5. Session Info

```
sessionInfo()
```

```
## R version 3.2.2 (2015-08-14)
## Platform: x86_64-pc-linux-gnu (64-bit)
## Running under: Ubuntu precise (12.04.5 LTS)
## 
## locale:
##  [1] LC_CTYPE=en_US.UTF-8       LC_NUMERIC=C              
##  [3] LC_TIME=en_US.UTF-8        LC_COLLATE=en_US.UTF-8    
##  [5] LC_MONETARY=en_US.UTF-8    LC_MESSAGES=en_US.UTF-8   
##  [7] LC_PAPER=en_US.UTF-8       LC_NAME=C                 
##  [9] LC_ADDRESS=C               LC_TELEPHONE=C            
## [11] LC_MEASUREMENT=en_US.UTF-8 LC_IDENTIFICATION=C       
## 
## attached base packages:
## [1] stats     graphics  grDevices utils     datasets  methods   base     
## 
## other attached packages:
## [1] knitr_1.17
## 
## loaded via a namespace (and not attached):
##  [1] backports_1.0.5 magrittr_1.5    rprojroot_1.2   tools_3.2.2    
##  [5] htmltools_0.3.5 yaml_2.1.14     Rcpp_0.12.9     stringi_1.1.2  
##  [9] rmarkdown_1.8   stringr_1.2.0   digest_0.6.12   evaluate_0.10
```
